# Network and synaptic mechanisms underlying high frequency oscillations in the rat and cat olfactory bulb under ketamine-xylazine anesthesia

**DOI:** 10.1101/2020.07.23.217604

**Authors:** Władysław Średniawa, Jacek Wróbel, Ewa Kublik, Daniel Krzysztof Wójcik, Miles Adrian Whittington, Mark Jeremy Hunt

**Affiliations:** Nencki Institute of Experimental Biology, 3 Pasteur Street, 02-093 Warsaw, Poland; University of Warsaw, Faculty of Biology, Miecznikowa 1, 02-096, Warsaw, Poland; Jagiellonian University, Faculty of Management and Social Communication, Jagiellonian University, 30-348 Cracow, Poland; University of York, Heslington, York, YO10 5DD, United Kingdom

**Author notes:** Correspondence: Mark Jeremy Hunt, Nencki Institute of Experimental Biology, 3 Pasteur Street, 02-093 Warsaw, Poland.

## Abstract

High frequency oscillations (HFO) are receiving increased attention for their role in health and disease. Ketamine-dependent HFO have been identified in cortical and subcortical regions in rodents, however, the mechanisms underlying their generation and whether they occur in higher mammals is unclear. Here, we show under ketamine-xylazine anesthesia, classical gamma oscillations diminish and a prominent > 80 Hz oscillation emerges in the olfactory bulb of rats and cats. In cats negligible HFO was observed in the thamalus and visual cortex indicating the OB was a suitable site for further investigation. Simultaneous local field potential and thermocouple recordings demonstrated HFO was dependent on nasal airflow. Silicon probe mapping studies spanning almost the entire dorsal ventral aspect of the OB revealed this rhythm was strongest in ventral areas of the bulb and associated with microcurrent sources about the mitral layer. Pharmacological microinfusion studies revealed HFO was dependent on excitatory-inhibitory synaptic activity, but not gap junctions. Finally, we showed HFO was preserved despite surgical removal of the piriform cortex. We conclude that ketamine-dependent HFO in the OB are driven by nasal airflow and local dendrodendritic interactions. The relevance of our findings to ketamine’s model of psychosis in awake state are also discussed.

## 1 Introduction

Local field potential (LFP) oscillations reflect synchronous activity of neuronal assemblies and are thought to play a crucial role in information processing [Engel and Singer, 2001, Colgin and Moser, 2010]. Recent years have witnessed a surge of interest in high frequency oscillations (HFO), also known as ripples, considered important for their roles in health and disease [Buzsáki et al., 2012, Khodagholy et al., 2017]. HFO have been investigated most notably in the rodent hippocampus, but are also found in diverse cortical, olfactory and limbic areas [Haufler and Pare, 2014, Zhong et al., 2017, Vaz et al., 2020], and have been linked to near-death states [Borjigin et al., 2013], seizures [Zijlmans et al., 2012], Parkinson’s disease [Foffani et al., 2003], and models of psychoses [Hunt et al., 2006]. Collectively, these results point to regional and state dependent mechanisms underlying HFO generation in the brain. N-methyl-D-aspartate receptors (NMDAR) are expressed widely in the CNS, and are well-known to be involved in learning and memory processes [Shimizu et al., 2000]. NMDAR antagonists like ketamine or phencyclidine, induce short-lasting psychotic states and is used to study synaptic mechanisms of psychoses in animal models [Frohlich and Van Horn, 2014]. To date, many groups, including our own, have shown that low-doses of ketamine (or related NMDA receptor antagonists) produce HFO in many cortical and subcortical areas [Hunt et al., 2006, Phillips et al., 2012, Caixeta et al., 2013, Flores et al., 2015, Kealy et al., 2017, Amat-Foraster et al., 2019]. However, there are important gaps in understanding ketamine-dependent HFO in the brain, most notably, what are the network and synaptic mechanisms underlying this activity, and does it occur in higher mammals.

Glutamate is the major excitatory neurotransmitter in the OB, and its NMDA receptors are expressed by the major cell types [Nagayama et al., 2014]. NMDA receptors are important at reciprocal dendrodendritic mitral-granule cell synapses, considered to underlie the generation of fast rhythms in the OB [Neville and Haberly, 2003, Fourcaud-Trocmé et al., 2014, Osinski and Kay, 2016]. We showed recently that the olfactory bulb (OB) is a strong generator of ketamine-dependent HFO in the brain of rodents [Hunt et al., 2019]. The OB appears to have a particularly privileged position since it can impose both fast and slow oscillatory activity in distant regions [Ito et al., 2014]. Although ketamine-dependent HFO has been well-documented, little is known about its mechanisms of generation or its occurrence in higher mammals. The OB is the first relay station of the olfactory system and receives direct sensory input from the olfactory nerve. Since their discovery in 1942, fast oscillations in the OB [Adrian, 1942], in particular gamma, have been the subject of growing investigations in vivo, in vitro and in modelling studies [Rojas-Líbano et al., 2014]. Indeed, it is well elucidated that NMDA receptors at reciprocal mitral–granule cell dendrodendritic synapses [Schoppa et al., 1998, Isaacson and Strowbridge, 1998, Chen et al., 2000] underlie gamma rhythmogenesis [Osinski and Kay, 2016]. Even in the absence of odors, in humans and rodents nasal respiration can powerfully entrain fast and slow oscillations in the OB, and other brain regions [Grosmaitre et al., 2007, Zelano et al., 2016, Zhong et al., 2017], for example delta oscillations during rodent anesthesia and quiet waking. Anesthetized states offer distinct advantages when studying fundamental neuronal processing, since core networks and respiration are spared. Although anesthesia is usually associated with reduced power [Bagur et al., 2018], and frequency [Ylinen et al., 1995] of fast oscillations, there is some evidence that HFO can be observed under ketamine-xylazine anesthesia [Grenier et al., 2001, Chery et al., 2014]. This provides an opportunity to probe the mechanisms of ketamine-dependent HFO rhythmogenesis in a straightforward manner devoid of behavioural confounds.

In this study, we found ketamine-xylazine anesthesia in rats and cats was associated with a switch from gamma activity and emergence of HFO in the OB. We found ketamine-xylazine HFO resembled the ketamine-dependent HFO found in the wake-related state, although of slower frequency. This rhythm was strongest in ventral parts of the OB, entrained by nasal respiration, and mediated by excitatory and inhibitory synaptic transmission. It was preserved despite removal of the piriform cortex indicating HFO arises through intrinsic OB circuitry. We conclude that ketamine-dependent HFO is a fundamental brain rhythm in mammals. Given the reduction in frequency we observed previous reports on classical gamma in some ketamine-xylazine anesthetised studies may have inadvertently detected ketamine-dependent HFO.

## 2 Results

### 2.1 Ketamine-xylazine and subanesthetic ketamine differentially affect gamma (30–65 Hz) and faster rhythms

We compared fast oscillatory activity recorded from LFP’s in the OB after a subanesthetic dose of ketamine 25 mg/kg and under KX anesthesia (ketamine 100 mg/kg + xylazine 10 mg/kg) and ketamine 200 mg/kg anesthesia (n=9 rats) (Fig. 1).

**Figure 1.**
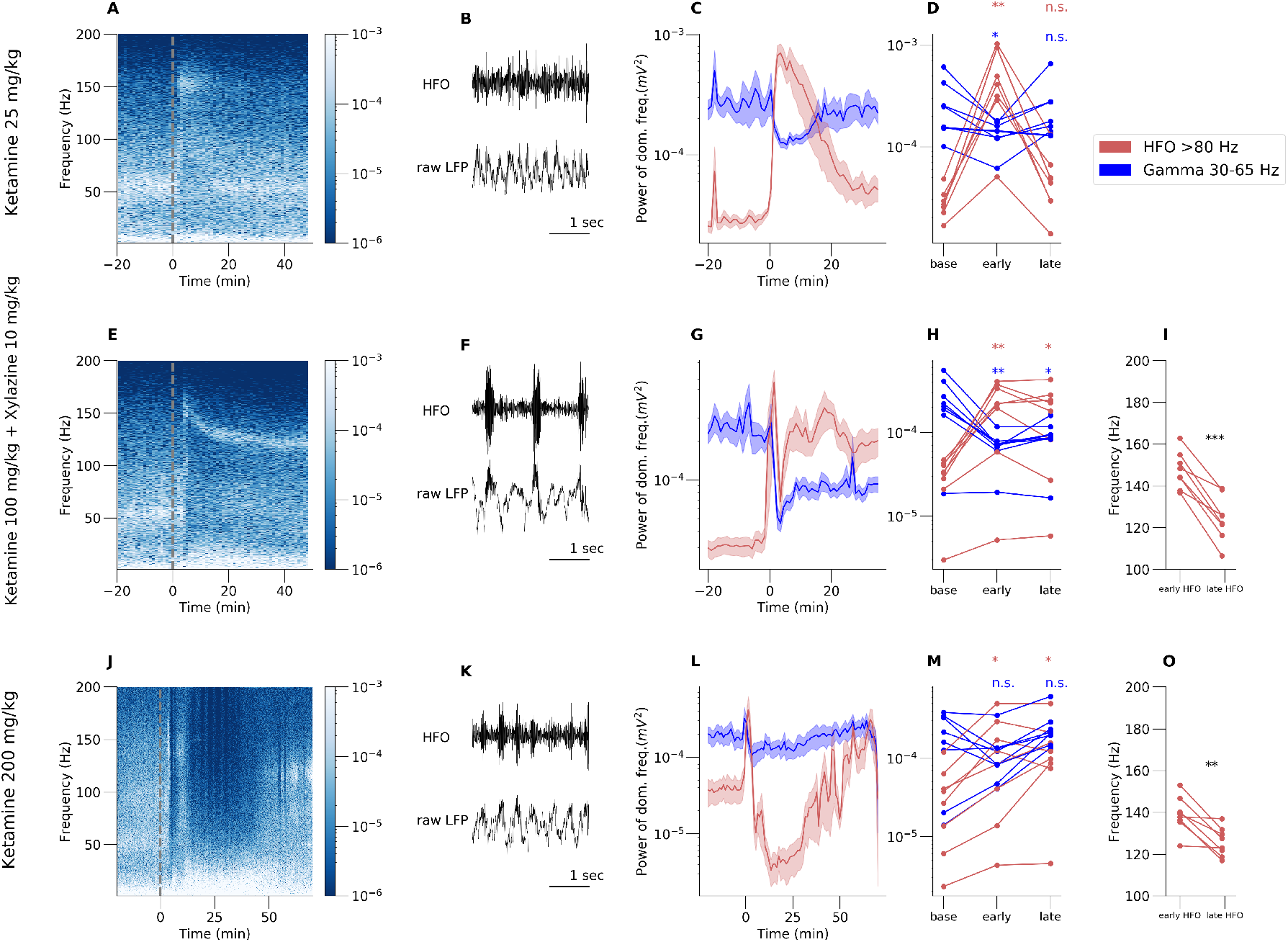
First row (A–D) presents results for subanesthetic ketamine 25 mg/kg, second row (E–I) for ketamine 100 mg/kg + xylazine 10 mg/kg (KX) anesthesia and third row for anesthetic dose of ketamine 200 mg/kg. First column (A, E and J): Example spectrograms computed from the olfactory bulb before and after injection. Dotted line at *t* = 0 is injection time. Second column (B, F and K): Example raw waveform (bottom) and filtered signal in HFO band 80-180 Hz (top). Third column (C,G and L): Extracted data from spectrogram of the power of dominant frequency in gamma (30–65 Hz) and HFO (80–180 Hz) range (average across rats n=9). Both subanesthetic ketamine and KX administration reduced gamma power and substantially increased HFO power. Third column (D,H and M): Individual traces of HFO and gamma power taken from three time points (mean of 5 min) 15 min before, 1 min after, and 50 (D and H) or 80 min (M) after injection. I and O: Comparison of the HFO frequency just after injection and at the end of the recording. Note the gradual reduction of frequency from 150 to around 120 Hz under KX anesthesia.

Immediately after injection of 25 mg/kg ketamine we observed substantial fast activity at 152.37±10.21 Hz and a concomitant reduction in power of 30–65 Hz activity (Fig. 1 A–C, early HFO p=0.0042, early gamma p=0.049, paired t-test, n=8). KX anesthesia, confirmed by loss of tail-pinch reflex and eyeblink reflexes, was also associated with initial fast activity (Fig. 1 E–H, early HFO p=0.0046, early gamma p=0.0077, paired t-test, n=8), which over the course of an hour gradually slowed in frequency to 124.43 ± 10.07 Hz (Fig. 1 I, p< 10^−4^, paired t-test, n=8). Like 25mg/kg ketamine, KX anesthesia also produced a concomitant reduction in 30–65 Hz activity (Fig. 1 E–H). An anesthetic dose of ketamine alone was associated with a reduction in fast activity which lasted around 45-60 min followed by emergence of HFO during the recovery phase, typically associated with recovery of the righting reflex (Fig. 1 E–H, late HFO p=0.023, late gamma p=0.32, paired t-test, n=8). HFO that emerge during recovery of the was slower 125.83 ± 8.66 in frequency than initial burst after injection (Fig. 1 O, p=0.0029, paired t-test, n=8). Attenuation of most EEG oscillatory activity (described as EEG holes) following an anesthetic dose of ketamine has been reported recently in sheep [Nicol and Morton, 2020] and we have observed previously that anesthetic ketamine attenuates HFO in the nucleus accumbens of rodents [Hunt et al., 2006].

We noticed that both fast oscillations after subanesthetic ketamine and KX anesthesia occurred as discrete bursts, lasting around 100 ms, and were nested towards the peaks of slow frequencies (Fig. 1 B,F and K). In rats injected with 25 mg/kg ketamine coupling occurred at theta frequencies, whereas in KX coupling occurred at slower delta frequencies.

In the same group of rats we injected xylazine 10 mg/kg in 4 rats and we observed no effect on 80–180 Hz activity (Fig. S1 A and B), and a trend for increased 30–65 gamma power was observed. This indicated that ketamine rather than xylazine was responsible for the effect on the high frequencies we observed.

### 2.2 Ketamine-xylazine HFO oscillations are present in the OB, but not thalamus or visual cortex in cats

A fundamental issue when examining under-reported brain activity is whether electrophysiological signatures are present in a higher mammals. We therefore extended our studies to include cats (n=3) and recorded simultaneously from the olfactory bulb, the lateral geniculate nucleus (LGN) of the thalamus, and from the visual cortex (Fig. 2 A1–A3). In OB of two cats ~ 90 Hz oscillation was recorded under KX anesthesia (Fig. 2 B2–B3). In the third cat histology revealed electrode placement at the edge of the OB which was associated with fast activity of comparable frequency but much smaller power (Fig. 2 B1). Consistent with our findings from rats, this fast activity occurred as discrete bursts (Fig. 2 A1). Importantly, HFO activity was not present in thalamus or visual cortices, demonstrating certain neuroanataomical selectivity to the OB under KX anesthesia. However, note that in awake rats, where the power of fast activity is higher, subanesthetic ketamine can induce HFO in many subcortical areas [Hunt et al., 2006, 2009]. In line with our previous findings that ketamine-related fast oscillations can be attenuated by various types of anesthesia [Hunt et al., 2009], here, we found that propofol anesthesia also attenuated fast oscillations associated with KX (Fig. 2 B2 and B3 insert). We also calculated the modulation index score (Fig. 2 C1–C3) for OB channels in each cat, see methods for computational details. Results strongly indicate that KX HFO in cats is coupled to local slow oscillations in OB (strong blue spot around 1 Hz-90 Hz pixel).

**Figure 2.**
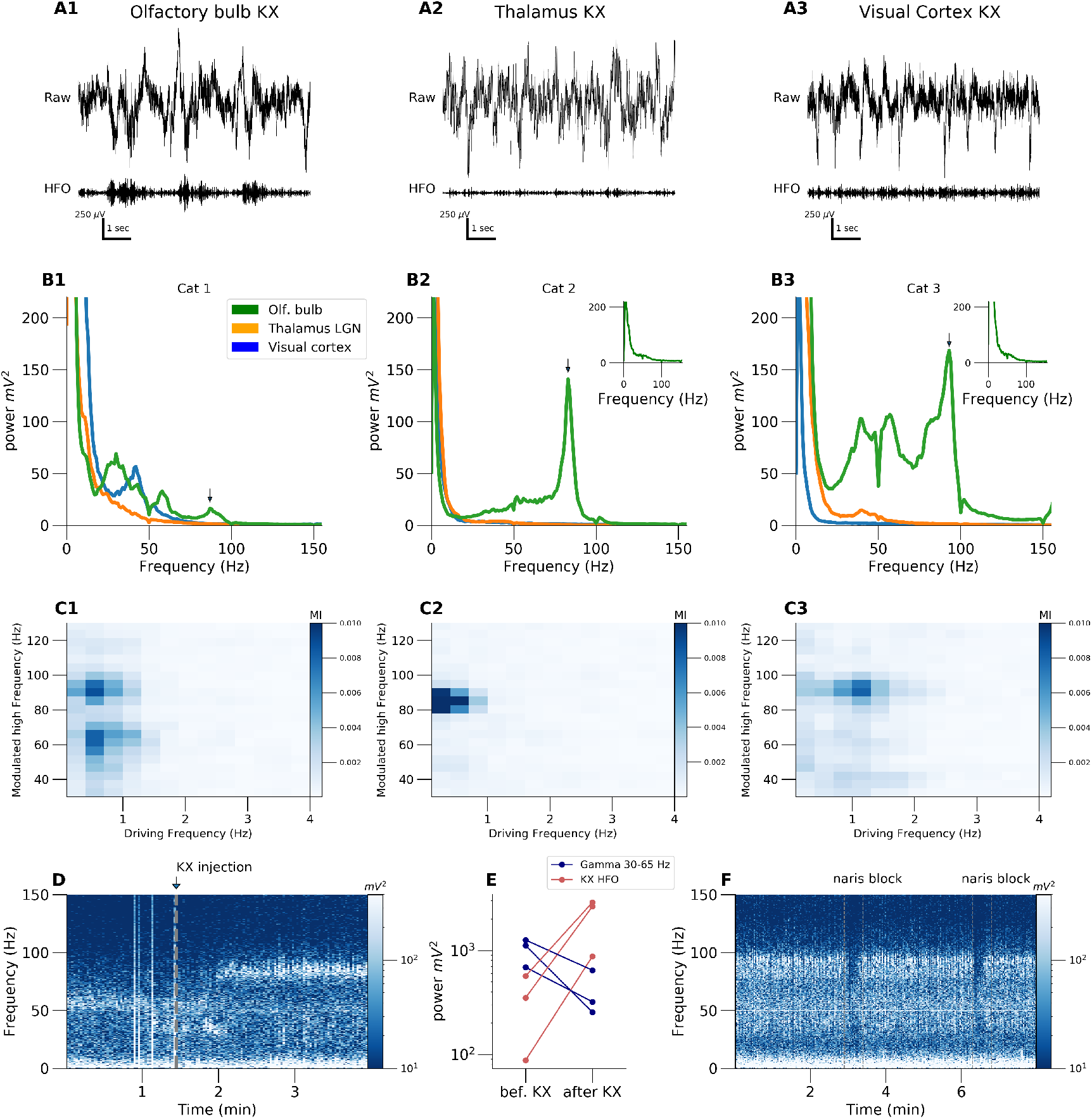
A: Example raw waveforms from one cat from three different brain regions. Raw signal is presented in top rows and 80–130 filtered signal is in bottom row. B: Power spectrum from 10 min of recordings. We observed ~ 90 Hz HFO separate band in all the recorded cats. In the cat with the electrode in posterior OB HFO was smaller. There was no activity of this type in other simultaneous recorded structures (Thalamus, Visual Cortex) nor in OB under propofol anesthesia condition (insets). C: Modulation index score computed from single OB channels for all three cats. Color strength of the ‘pixel’ represents power of the modulation for a given slow (driving frequency) and fast (modulated frequency) oscillation extracted from the raw signal. D and E show transition from low ketamine-xylazine to higher dose. We reported increase in 90 Hz band with parallel decrease of the power of gamma activity. The same phenomena were observed in rats; compare Fig. 1. We checked what happens if we unilaterally block naris of the cat and we saw rapid reduction in HFO power (F) but not 160 Hz activity of unknown origin (supplementary figure).

KX was infused intravenously at a rate of 0.2 ml every 20 min which provided a window to determine any electrophysiological changes just after administration of KX. Prior to administration gamma ~ 60 Hz oscillations were present in the OB, then immediately after infusion a clear 90 Hz oscillation was visible (Fig. 2 D). This effect was reproducible across different infusions (Fig. 2 E). Nasal respiration is known to drive fast and slow oscillations in the OB [Neville and Haberly, 2003]. To test if the same held true for KX fast oscillations we also applied short unilateral naris blockade, in two cats, and found this was associated with a reduction in 90 Hz oscillation power (Fig. 2 F). In one cat, in addition to the 90 Hz activity (described above) we also observed a strong ~ 160 Hz oscillation(Fig. S2 A and B) present exclusively in the bulb. We do not understand the origin of this oscillation which was not responsive to unilateral naris block (Fig. S2 C and D).

### 2.3 Ketamine-xylazine dependent high frequency oscillations are entrained by nasal respiration

Slow oscillations in the mammalian OB have long been known to be closely related to nasal respiration. We investigated the relationship between nasal respiration and KX fast oscillations further using thermocouples implanted in the naris of rats. These KX experiments, and those described hereafter, were carried out after initial isoflurane anesthesia for surgical procedures. As expected, shortly after injection of KX, fast rhythmic activity was visible in the raw LFP, nested on a slower oscillatory rhythm (Fig. 3D and E). However, prior isoflurane exposure reduced mean frequency of KX-HFO compared to KX alone (Fig. S3B, p=0.0028, paired one-way ANOVA)). For this reason, in these experiments the 80–130 Hz band was used. Although gamma oscillations in the OB can reach 80 Hz, considering that gamma power is largely attenuated under KX anesthesia, there would be little gamma contamination.

**Figure 3.**
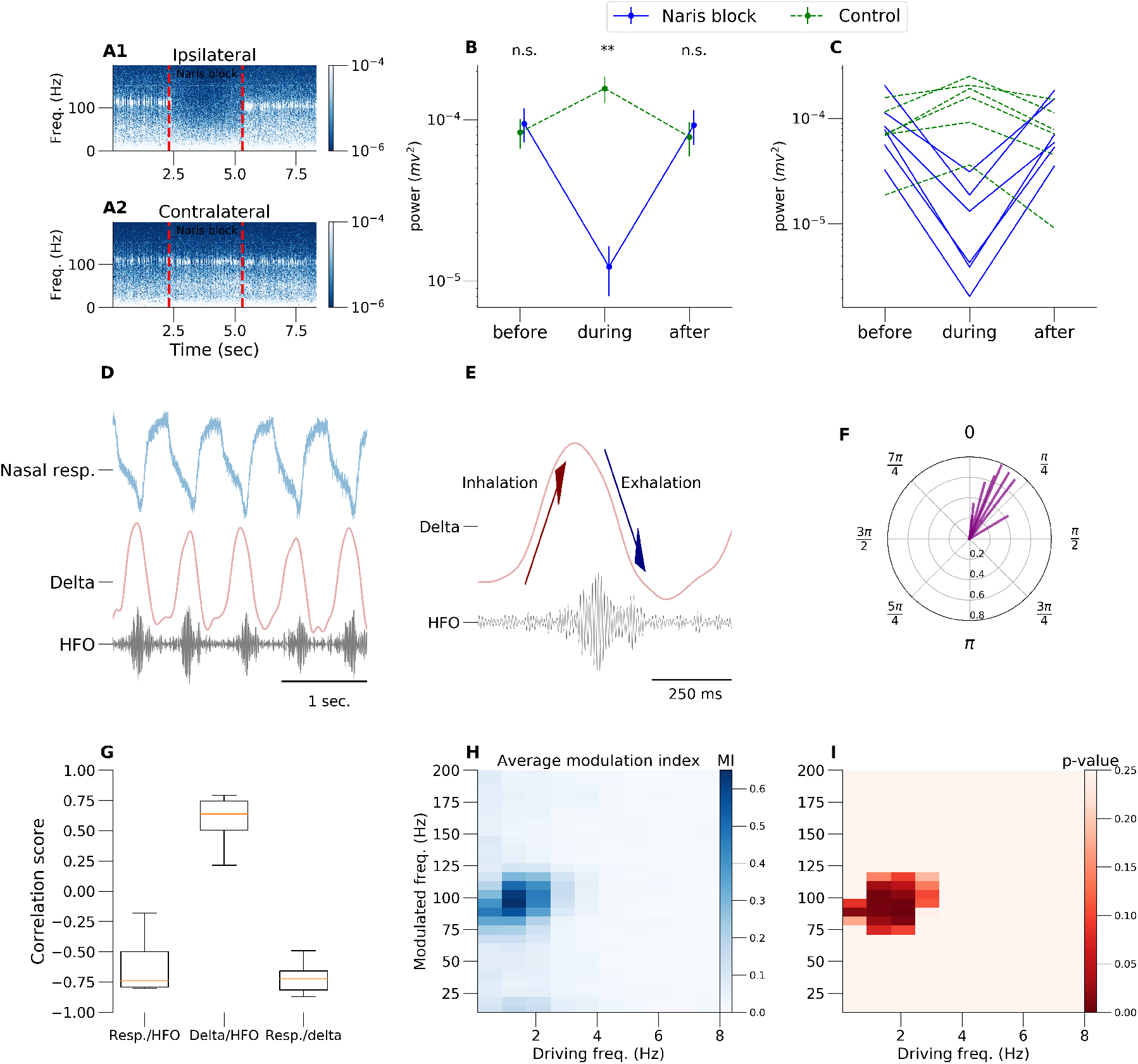
All presented results are during ketamine-xylazine (KX) anesthesia. A: Spectrogram computed from ipsilateral and contralateral to the naris blockade OB channels. Dotted lines relate to naris blockade time. B: Extracted data from spectrogram of power of dominant frequency in HFO 80–130 Hz range (average across rats n=8). Unilateral naris blockade reduced HFO power on the ipsilateral side but surprisingly also increased HFO power contralaterally to the blockade. Traces from individual rats are shown in C. D: Simultaneous OB LFP and respiratory rhythm recorded using thermocouples. E: Example recordings of simultaneous OB LFP for delta (0.3–3 Hz) and HFO (80–130 Hz) with arrows showing breath cycle phase. F: Phase plot of the peak of the burst of KX HFO relative to delta peak (phase zero). Each radius represents ITPC (see methods) of one rat. G: Waveform correlation scores calculated for 5 min for respiration rhythm and OB LFP delta (first box plot), delta and the OB LFP HFO envelope (second box plot), and respiration rhythm and OB LFP delta (third box plot). Box plots show first to third quartile of the correlation scores, median (orange line) and all data range of n=8 rats (error bars). H: Modulation index matrix (averaged across rats n=8) for 5 min of continuous LFP signal under KX anesthesia. I: Resampling (across rats) statistic for modulation index “pixels”. Control group for resampling are rats under isoflurane anesthesia.

Consistent with our findings from cats, unilateral naris blockade immediately reduced the power of 80–130 Hz activity (Fig. 3A–C, p=0.0013, one-way ANOVA). The reduction of 80–130 Hz activity power in the OB occurred exclusively on the ipsilateral side and quickly recovered when blockade was removed. Interestingly, a small increase in 80–130 Hz activity power was recorded in the contralateral side.

Fig. 3D shows an example recording of nasal respiration, and its relation to < 2Hz, and 80–130 Hz activities recorded in the OB. Note that bursts temporally correlate with peaks of the local delta oscillation and troughs from thermocouple signal (Fig. 3D and G). We computed correlation coefficients between nasal respiration, <3 Hz filtered signal and the 80–130 Hz envelope (Fig. 3G). Interestingly, phase analysis (Fig. 3 E and F) showed that the peak of the envelope of fast oscillatory bursts were time-locked to slower oscillations on the descending phase for KX (Fig. 3 F). The peak phase of local slow LFP rhythm corresponds to inhalation-exhalation transition in breathing cycle (Fig. 3E). We also calculated the modulation index score (Fig. 3H and I), see methods for computational details, which confirmed that 80–130 Hz oscillations are driven by nasal respiration.

### 2.4 80–130 Hz oscillations are associated with dipole-like current sources around the mitral cell layer

To determine if 80–130 Hz activity within the bulb was localised to particular layers we mapped 80–130 Hz activity in the OB using 32 site linear silicon probes of two densities (3.2 mm long with 100 *μ*m spacing and 0.64 mm long with 20 *μ*m spacing) (Fig. 4 A and E). 80–130 Hz oscillations were largely coherent across the OB including contacts up to 3.2 mm apart, the greatest distance we measured. Notably, 80–130 Hz power was substantially larger deep in the granule layer which was clearly visible in the 3.2 mm probe recordings (Fig. 4 B). We observed a sharp reduction in power close to the mitral layer (Fig. 4 C and G) where 80–130 Hz activity shifted in phase (Fig. 4D, F and H). For electrodes that did not cross the mitral layer 80–130 Hz oscillations were synchronous across all contacts, on a cycle by cycle basis. We found that the slow 2Hz oscillation also reversed phase, but more ventrally and within the EPL/glomerular layers (Fig. 4 D and H). Since field potentials reflect neuronal activity from a broad area we next reconstructed the underlying sources of 80–130 Hz oscillations and the 2 Hz oscillation. We used the kCSD method to accurately identify the spatial and temporal profiles of changes in membrane current, underlying the field potential (see Methods for computational details). In the raw signal, triggered on 80–130 Hz oscillatory events, we found dipole-like spatiotemporal profiles across the mitral and granule cell layers (Fig. 4 I). CSD reconstruction (average for N=8) was filtered for slow frequencies (0.3–8 Hz) and we observed dipole-like structure that propagates from glomerular layer, before emergence of 80–130 Hz activity, to EPL layer as the 80–130 Hz power rises in time (Fig. 4 J). We next filtered the CSD in the 80–180 Hz band and found strong dipoles around the mitral layer (Fig. 4 K average across 8 rats). Individual traces from time point zero, from all rats, showing delta and 80–130 Hz dipoles are presented in supplementary Fig. S4, respectively. Analysis of the Multi Unit Activity (MUA >500 Hz LFP oscillations) shows that 80–130 Hz oscillations are driven by local spiking of the mitral/tufted cells (Fig. 4 L). Correlation computed between local 80–130 Hz activity and envelope of MUA density (see methods) shows that spikes does not change phase together with 80–130 Hz — spikes occur in a trough of the oscillation around EPL/mitral cell layer.

**Figure 4.**
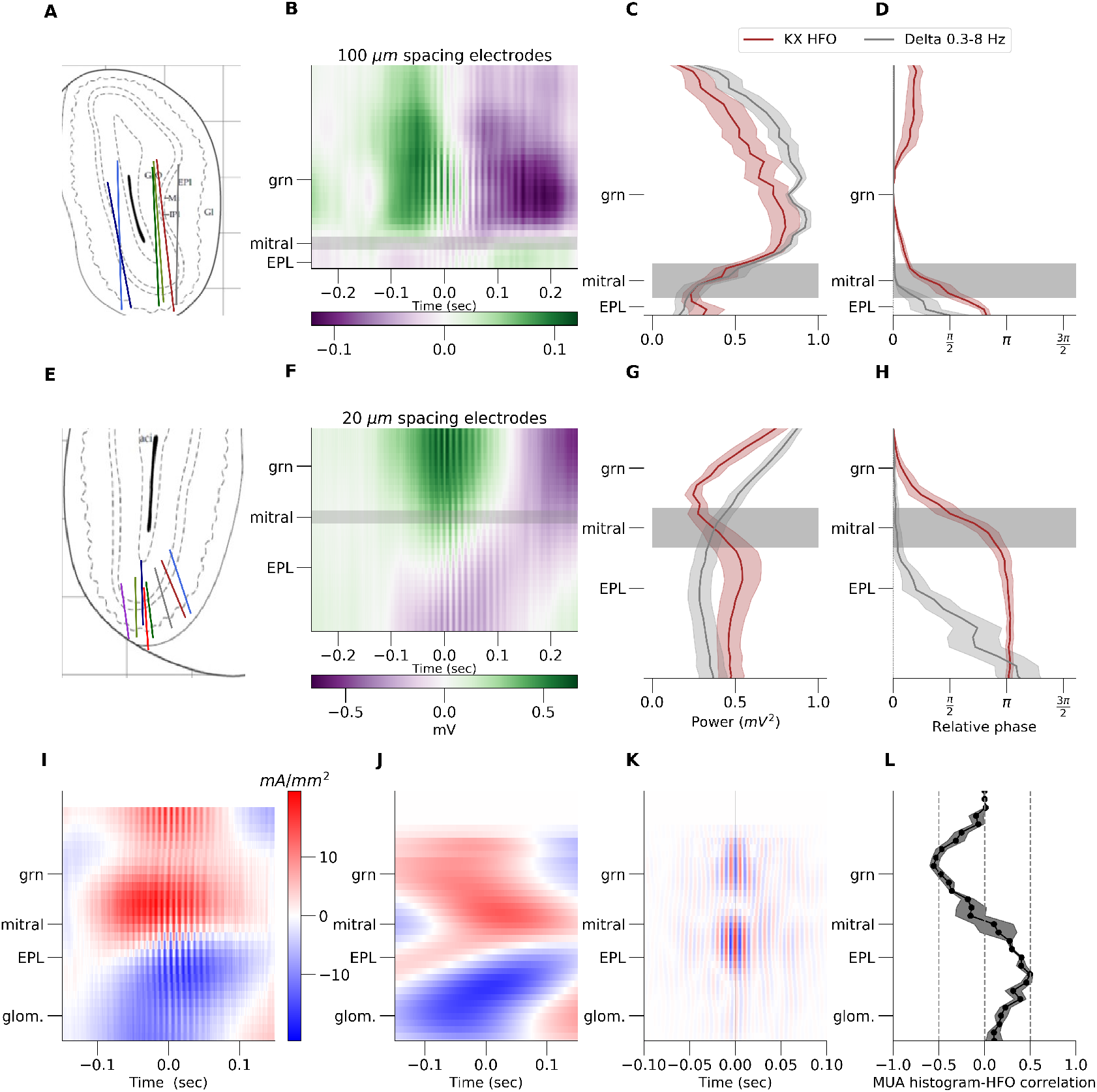
Results presented in A to D corresponds to 3.2 mm electrode with 100 *μ*m inter-electrode distance. A: Histology illustration based on n=6 rats OB stainings. B: Example raw LFP from 32 channels (y axis). Colorbar scale is in mV. C: Power of HFO (80 – 180 Hz) from 32 channels. Power decreases around mitral layer. x-scale is in mV ^2^ D: Phase of HFO (80 – 130 Hz) – HFO changes phase of oscillation around mitral layer, Delta changes below mitral. x-scale is in degrees. Results presented in E to H corresponds to 0.64 mm electrode with 20 um inter-electrode distance and shows the same type of analysis as in A-D (n=8). Results presented from I to K shows CSD analysis for 20 *μ*m spacing electrode. I: raw example CSD. J: CSD filtered in 0.3-3 Hz band. Plot were then aligned in respect to the histology and averaged across rats n=8. K: CSD filtered in 0.3-3 Hz band. L: Average correlation plot between of local HFO and MUA density (n=4).

### 2.5 Blockade of GABA-A and AMPA but not gap junctions disrupts 80–130 Hz activity

CSD and MUA analyses strongly suggest that mitral/tufted cells are involved in the generation of 80–130 Hz rhythm, consistent with our finding after ketamine in freely moving rats [Hunt et al., 2019]. Excitation, inhibition and gap junctions can all underlie mechanisms of fast oscillatory activity in the brain [Grenier et al., 2001]. To unveil the receptor mechanisms generating 80–130 Hz under KX, we carried out a series of pharmacological experiments using unilateral infusion of bicuculline, NBQX, or carbenoxolone (0.5 *μ*l) (Fig. 5). In the presence of KX, the effect of bicuculline was rapid with potent reductions in 80–130 Hz power being visible in many cases during the infusion (sal. vs bic. n=5, p=0.0053, one-way ANOVA). By contrast, slow delta oscillations in the OB, largely dependent on respiration, remained clear in the raw signal up to 20 min post infusion and the absolute power of delta was not affected by drug infusion (Fig. 5A–F) (sal. vs bic. n=5, p=0.098, one-way ANOVA). NBQX infusion also reduced 80–130 Hz power (sal. vs NBQX n=7, p=0.00011, one-way ANOVA), however, in about half the rats we observed a delayed reduction in the amplitude of slow oscillations but this was well after effects on 80–130 Hz activity occurred (saline vs NBQX n=7, p=0.012, one-way ANOVA). Thus, reductions in 80–130 Hz activity post NBQX were unlikely to be due to reduced sensory input, but rather due to AMPA blockade within the OB circuitry. These findings support a role for both GABA-A and AMPA receptors in the generation of the 80–130 Hz oscillation we recorded. Carbenoxolone did not affect 80–130 Hz activity (sal. vs carb. n=4, p=0.54, one-way ANOVA). Neither delta oscillations were affected (sal. vs carb. n=4, p=0.96, one-way ANOVA) (Fig. 5G–I). Thus, most probably, gap junction connections do not play a role in 80–130 Hz rhythm generation.

**Figure 5.**
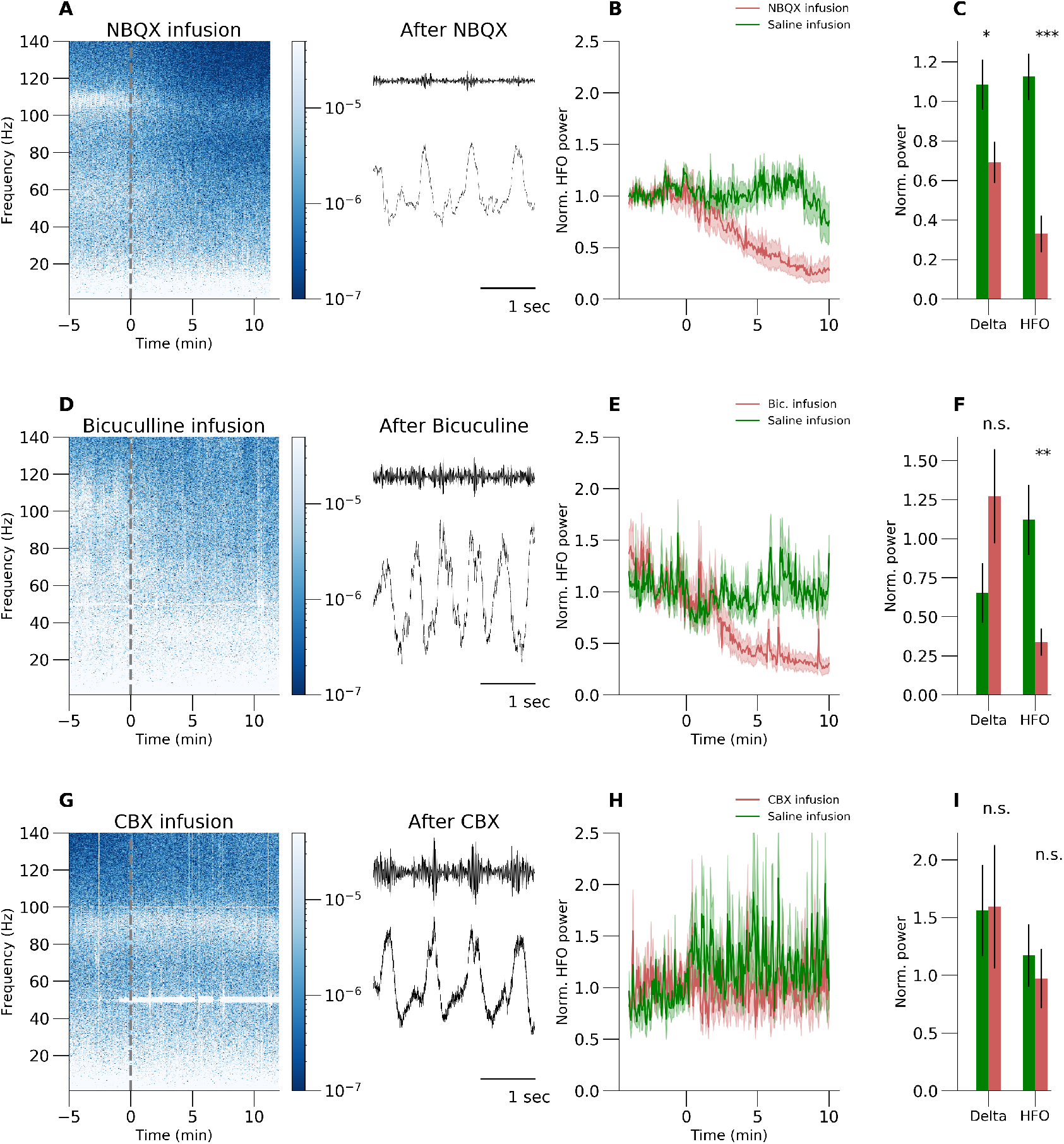
HFO under ketamine-xylazine are inhibition and excitatory-based but not dependent on gap junctions. A to C show results for NBQX local infusion to the bulb (red lines and red bars) which caused significant reduction in power of the HFO (B and C, n=7). Picture A is an example spectrogram and dotted line refers to the moment of the infusion. D to F and G to I show the same results and analysis as the top row. GABAA receptor blockade with bicuculline infusion (red lines and bars) caused the same effect — reduction in power of the HFO (red lines and bars in E and F, n=5). Example waveforms for a given drug after infusion are presented in the second column. However, gap junction blockage with carbenoxolone seems to have no effect on HFO (H and I, n=4).

### 2.6 Removal of piriform and surrounding regions do not disrupt 80–130 Hz KX oscillations

The piriform cortex is a major downstream structure for OB projections. Pyramidal neurons of the piriform cortex also send dense projection back to OB [Boyd et al., 2015, Diodato et al., 2016] which have been proposed to control the gain of OB activity [Boyd et al., 2012]. This prompted us to examine the potential role of the piriform cortex, and surrounding regions, on 80–130 Hz activity recorded in the OB. Under KX anesthesia, we gradually removed cortical tissue, starting laterally and moving medially. We did this until the base of the skull was exposed to ensure that a large amount of the piriform cortex has been extracted (Fig. 6A). Comparison of LFP oscillations from the dissected vs. intact side had similar power of 80–130 Hz activity (Fig. 6B–D, p=0.64 and p=0.37, paired t-test, n=9). There was neither significant change in frequency after brain removal (Fig. 6E, p=0.34 and p=0.55, paired t-test, n=9). We did not dissect right up to the midline due to the presence of major vessels. However, in four rats the anterior commissure (which carries centrifugal fibers to the OB) was also partially or fully transected.

**Figure 6.**
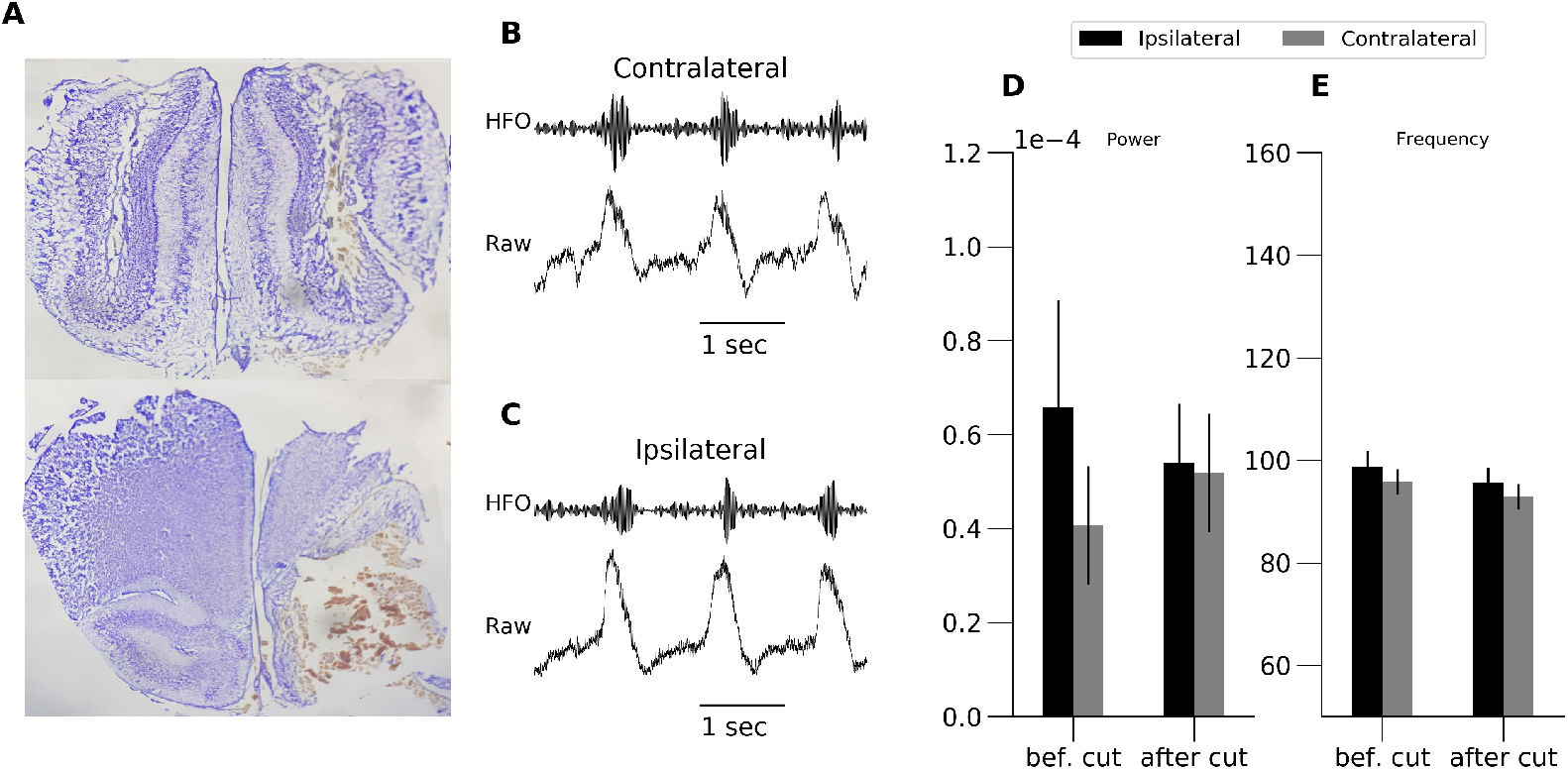
HFO do not depend on the piriform cortex or centrifugal input. Picture A shows example conceptual histology of rat in which we removed part of frontal lobe. B and C presents example waveforms at the contralateral and ipsilateral side after the cut was made. We did not see any differences after brain removal neither in power nor in frequency of the oscillations (D and E, n=9). This support the theory that HFO oscillations are generated locally in olfactory bulb by neuronal circuits.

We were not able to dissect to the midline due to the presence of the anterior rostral nerve, which is necessary for circulation of blood to the OB and other anterior brain regions. Unexpectedly, on occasions when this vessel was punctured we observed a transient increase in 80–130 Hz activity followed by attenuated activity. We do not currently understand this hemodynamic effect but suspect it is associated with anoxia, and appears in line with another study showing cardiovascular arrest also induced a transient burst of fast oscillatory activity [Borjigin et al., 2013].

## 3 Discussion

### Nasal respiration is critical for ketamine 80–130 Hz activity

Here we show that a 80–130 Hz rhythm emerges under KX anesthesia in the OB of rats and cats. This rhythm shows some degree of neuroanatomic specificity, being large in amplitude in the OB, relatively weak at the OB border, and absent in thalamic and visual cortices. 80–130 Hz activity occurs as discrete bursts frequently coupled to nasal respiration, indeed naris occlusion attenuates this rhythm. The olfactory bulb receives direct input from the olfactory nerve and nasal respiration has long been known to entrain the rhythmic activity in olfactory networks [Adrian, 1942, Macrides and Chorover, 1972]. We found that KX mitral/tufted neurons fire in phase with 80–130 Hz, consistent with a previous finding under KX reporting mitral/tufted firing over 100 Hz and associated with inhalation/exhalation transitions [Burton and Urban, 2014]. Our previous work in freely moving rats given a subanesthetic dose of ketamine also showed spiking phase locked to a fast 130–180 Hz activity [Hunt et al., 2019]. Nasal respiratory therefore provides a gate for the firing of OB projection neurons [Wachowiak, 2011] and the emergence of ketamine-dependent fast rhythms. Using a 3.2 mm probe we mapped, almost entirely, the dorsal-ventral aspect of the rat OB. The 80–130 Hz rhythm was progressively larger in amplitude in mid-ventral areas of the OB. Axonal projections of mitral/tufted cells are highly spatially localized, whereby axons of cells in the ventral OB travel through ventral regions of the granule layer to the olfactory tract [Hintiryan et al., 2012, Fukunaga et al., 2012]. CSD analyses, revealed the presence of two spatially and temporally distinct current dipoles. A sink, around 1 Hz, followed the breathing cycle, was localized close to the EPL layer and a faster 80–130 Hz microcurrent dipole was found closer to the mitral layer. Current sinks indicate flow of the positive ions (i.e., potassium, calcium) into neurons [Nicholson and Freeman, 1975, Mitzdorf, 1985]. Stimulation of the olfactory nerve to mimic afferent input, generates a sink in the glomerular and EPL layers [Neville and Haberly, 2003, Kay, 2014] and induces slow spontaneous 0.5–5 Hz rhythms [Fourcaud-Trocmé et al., 2014] consistent with respiratory drive.

The OB receives excitatory and modulatory input from the piriform cortex and centrifugal fibers [Nunez-Parra et al., 2013, Boyd et al., 2015]. However, these projections do not appear to make a significant contribution to the 80–130 Hz rhythm since it persisted even when large amounts of the piriform and commissural areas were removed. Notably, slow respiratory rhythms were also preserved confirming the primary drive arising from nasal input. These inputs are not essential for gamma oscillations in the OB [Neville and Haberly, 2003] indicating 80–130 Hz oscillations, like gamma, are generated within by the intrinsic circuitry of the OB.

### 3.1 Mechanisms of ketamine’s effect on fast oscillations

Both NMDA blockade and nasal respiration are prerequisites for the 80–130 Hz rhythm we recorded. Importantly, this activity was not present prior to NMDA blocker injection and was attenuated by isoflurane (rats) and propofol (cats) anesthesia. Xylazine alone had no significant impact on 80–130 Hz power indicating dependence of this rhythm on NMDA receptor blockade. Fast oscillations typically require interplay between excitatory and inhibitory transmission [Buzsáki et al., 2012]. Within the OB fast oscillations can be generated through glutamatergic release by mitral cell dendrites onto granule cell spines and reciprocal GABA release to locally inhibit depolarization spread [Chen et al., 2000]. Our microinfusion studies showed that inhibitory and excitatory components are required for the emergence of 80–130 Hz activity. By contrast, gap junctions, which can also underlie both physiological and pathological fast rhythmogenesis [Traub et al., 2002] and are expressed by mitral cells [Christie and Westbrook, 2006] were not implicated in this rhythm. Gamma can be evoked *in vitro* in OB slices demonstrating the intrinsic nature of this rhythm [Lagier et al., 2004, Galán et al., 2006]. Although higher frequencies, such as ripples (100–200 Hz), have been observed in hippocampal and cortical slices [Draguhn et al., 1998, Traub et al., 2013] to our knowledge there are no reports of frequencies in this band in OB slices.

Although oscillations are often investigated in single isolated bands, important inter-band interactions can underlie rhythmogenesis in the brain. In our rat and cat studies we observed a rapid switch from classical 30–70 Hz gamma to 80–130 Hz activity. Modeling studies have shown that granule cell excitability can control the frequency of OB network oscillations, and account for switches in state from beta to gamma oscillations [Osinski et al., 2018]. NMDA receptor antagonists preferentially target inhibitory interneurons [Li et al., 2002, Homayoun and Moghaddam, 2007] and optogenetic silencing of granule cells can also reduce gamma oscillations in the OB [Fukunaga et al., 2014] mimicking the ketamine effect. There is also evidence that NMDAR antagonists, including ketamine, can disinhibit granule cell-mediated inhibition of mitral cells [Wilson et al., 1996]. Given the above it is plausible that ketamine-induced reduction in gamma power are due to direct disruption of granule-mitral interactions.

NMDA blockade would reduce excitatory drive at mitral-granule cell synapses. Reciprocal GC-MC communication would be suppressed since NMDA receptors are necessary for Ca influx and subsequent GABA release [Chen et al., 2000]. Thus other excitatory-inhibitory networks would be predicted to underlie the generation of ketamine-dependent fast rhythms. There is an abundance of other types of inhibitory interneurons within the OB which can shape M/T behavior [Burton, 2017]. How might such fast oscillations be achieved in the OB? Clues may be found in other areas known to generate both gamma and faster oscillations. A good example is the hippocampus, where inhibitory PV-positive basket cells generate a ripple frequency. PV positive interneurons have been identified in the EPL layer of the OB, which make reciprocal contacts the perisomatic region of mitral cells [Kosaka et al., 1994]. They have morphological similarities to basket cells [Crespo et al., 2013]. In the OB, as elsewhere in the brain, PV interneurons are fast spiking [Mountoufaris et al., 2017]. In the OB, these cells can fire around 170 Hz, and are modulated by respiration [Kato et al., 2013]. They are thought to mediate broad lateral inhibition since they receive input and inhibit MC along their long secondary dendrites [Miyamichi et al., 2013] Importantly, excitation of PV-positive interneurons by mitral cells is chiefly mediated by AMPA receptors, with only a weak contribution by NMDA receptors [Kato et al., 2013]. Although PV-positive cells innervate mitral cells more densely than granule cells, very little is known about the capacity of this network to generate oscillations, however, given the precedent that these cells can generate fast rhythms (>100 Hz) we hypothesise that this circuit underlies the fast rhythm observed here [Kato et al., 2013]. We speculate that the movement of air across nasal mechanoreceptors stimulates the olfactory nerve, which in turn depolarizes MC via their dendrodendritic synapses, to induce burst firing in PV cells, which generates an E-I rhythm, which is turned off during expiration until the next breathing cycle. Reduced inhibitory tone at GC-MC synapses produced by NMDA blockade further increases the excitability of MC possibly enhancing this rhythm.

The OB also contains deep short axon interneurons which can modulate the firing of projection neurons, however, their firing frequencies would be unlikely to support fast rhythm [Burton et al., 2017]. Depolarization of mitral cells, expected to occur following disinhibition by granule cells, is associated with progressive increases in frequency and power of subthreshold oscillatory activity; however, since field activity is independent of excitatory and inhibitory transmission, and below 50 Hz, it would not account for the ketamine-dependent rhythm we observed [Desmaisons et al., 1999].

Given the above, we propose dynamical independent excitatory-inhibitory networks govern gamma (granule-mitral) and higher frequency (PV-mitral) oscillations in the OB. Differential sensitivities of these circuits to NMDA receptor blockade permit emergence or inhibition of fast brain rhythms.

### 3.2 Relevance to wake-related rhythms

The presence of 80–130 Hz activity in cats suggests this may be a fundamental effect of ketamine on mammalian olfactory networks. Fast oscillations produced by KX (80–130 Hz) and those occurring in freely moving rats after low-dose ketamine (130–180 Hz) were qualitatively similar. As shown in Figure 1 the 80–130 Hz rhythm closely paralleled the faster 130–180 Hz rhythms associated with subanesthetic ketamine in freely moving rodents. For example, 1) both oscillations occurred in bursts nested towards the peaks of slower rhythms; 2) both rhythms reversed phase in the vicinity of the mitral layer; 3) nasal airflow drives fast oscillations under KX (shown here) and subanesthetic ketamine in freely moving rats (submitted). Although anesthesia tends to attenuate most activity > 40 Hz there are some examples of ripple frequencies recorded under anesthesia; for example, Ylinen reported that under urethane and ketamine anesthesia the frequency of ripples was slower (100–120 Hz) than in the awake rat (180–200 Hz) [Ylinen et al., 1995]. Additionally, Neville and Haberly also reported that the frequency of discrete gamma and beta oscillatory bands is slower under urethane that the corresponding oscillations in awake rats [Neville and Haberly, 2003]. Finally, we have shown previously, the frequency of ketamine-dependent fast oscillations are highly dynamic and can drop by as much as 80 Hz after antipsychotic injection [Olszewski et al., 2013] and are likely to be related in awake and KX-anesthetised states. In summary, ketamine-dependent HFO is a highly dynamic band whose frequency varies according to state, and HFO recorded in awake and KX-anesthetized states appear to be related.

## Methods

### Surgery and chronic recordings

All experiments were conducted in accordance with the European community guidelines on directive 2010/63/UE on the protection of animals used for scientific purposes and approved by a local ethics committee. Thirty male Wistar rats (250–350 g) were used in this study. In 8 rats, tungsten electrodes (125 *μ*m, Science Products, Germany) were implanted bilaterally in OB (AP+7.5, ML=+0.5, DV=3–3.5 mm). LFP recordings were carried out before and after injection intraperitoneal of ketamine 25 mg/kg, and anesthetic doses of ketamine 100 mg/kg + xylazine 10 mg/kg (KX), and ketamine 200 mg/kg with 3–4 days separating each recording session (n=9). Under KX anesthesia the left or right naris was occluded using a soft piece of silicon rubber to block nasal respiration. At the end of the experiments the brains were post-fixed in 4% paraformaldehyde solution. Brains were dissected and placed in a 10% followed by 30% sucrose solution for 2–4 days. Electrode locations were determined on 40 *μ*m Cresyl violet (Sigma, UK) or Hoechst (Sigma, UK) stained sections.

### Thermocouple and LFP recordings

For acute studies, 8 rats were initially anesthetized using isoflurane during which time electrodes were implanted in the olfactory bulb and thermocouples in the frontal recess of the naris on the ipsilateral side. When electrodes were in place isoflurane anesthesia was replaced with KX by gradual injection of the cocktail and removal of isoflurane. Initial isoflurane exposure was necessary due to well-documented variable surgical plane anesthetic responses in rats compared to KX alone [Struck et al., 2011].

### Silicon probe recordings

A total of 14 rats were used for spatial mapping of oscillatory activity in the rodent OB. Rats were prepared for acute recordings as described above. Recordings were carried out in the OB using 32-channel silicon probes Edge-10mm-100-177 (N=6 rats, Neuronexus) and Edge-10mm-20-177 (N=8 rats). The electrodes were separated by an interelectrode distance of 100 (long probe) and 20 *μ*m (short probe). Prior to recording electrodes were dipped in a 5% solution of DIi (Sigma) dissolved in DMSO (Sigma). The track of the electrode was visualized using a fluorescent microscope.

### Local drug infusion

Rats were initially anesthetized using isoflurane and implanted with a guide-electrode complex in the left and right OB. Following implantation, isoflurane was substituted for KX anesthesia (see above for further information). LFPs were recorded bilaterally and when a stable fast oscillation was visible (80–130 Hz). NBQX (2 *μ*g, Sigma, n=7), Bicuculline methiodide (0.05 *μ*g, Sigma, n=5) and Carbenoxolone disodium salt (1 *μ*g, Sigma, n=4) were infused into the bulb. Saline was infused to the opposite bulb. For infusion, internal cannulae (28 gauge, Bilaney) that extended 2mm below the tip of the guides were inserted for 60s followed by 60s infusion of CNQX, bicuculline, CBX or saline (volume 0.5 *μ*l). Rats were recorded for 30 min post infusion and were then humanely killed using an overdose of anesthesia and their brains dissected for histological processing.

### Dissection of brain tissue

Rats were initially anesthetized by isoflurane for implantation of electrodes in the left and right OB. Following removal of the overlying skull isoflurane anesthesia was was replaced by KX. Under stable KX anesthesia we drilled the perimeter of a large cranial window approx. 7mm × 7mm (left hemisphere) from the midline to the lateral edge of the skull and removed the overlying bone. The exposed brain was dissected (using aspiration and a scalpel) in a lateral-medial direction until the base of the skull had been reached. LFPs from the left and right OB were recorded prior to and immediately after dissection.

### Cat experiments

All experiments were conducted in accordance with the European community guidelines on directive 2010/63/UE on the protection of animals used for scientific purposes and approved by a local ethics committee. Three healthy, mature cats (1 female, 2 male) were used. Cats were initially anesthetised using a bolus ketamine-xylazine i.p. injection . An intravenous catheter was implanted in a saphenous vein for supplementary ketamine-xylazine administration and 0.1-0.2 ml was administered every 20 min. Cat’s were placed in a stereotaxic frame and electrodes (32-channel silicon probes or in-house 16-channel electrodes made of 25 mm tungsten wire) implanted in the OB (AP+16,5, ML=−1,5, DV=3–4 mm), Thalamus nucleus (AP 6,5, ML=−9, DV=13,5 mm) and visual cortex (AP −16, ML=−1–9, DV=1–2 mm). Following acquisition of electrophysiological data the naris was briefly closed for 10 seconds (in 2 cats), anesthesia was then replaced with propofol for visual presentation experiments (not presented here).

### Data analysis

Scripts and files for figures generation are stored at: https://github.com/wsredniawa/KX

Recorded signals were processed using `SciPy signal` and `NumPy Python` libraries. Analysis included bandpass filtering using Butterworth filters. Power of dominant frequency and dominant frequency were evaluated using Welch transform from 60 seconds windows. To establish phase relation in KX HFO we first used Hilbert transform to find a maximum activity of HFO burst and then computed shift in time relative to peak of delta oscillations (score is rescaled to radians). Several hundred HFO bursts were used to compute intertrial phase clustering (ITCP) defined as 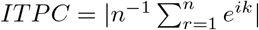, where *k* is the relative phase of the burst and *n* is the number of trial.

To study correlation of thermocouple’s rhythm and LFP signal oscillation we filtered the signal in delta frequencies 0.3-5 Hz and KX HFO 80–130 Hz. We computed a Pearson correlation score between the delta band of thermocouple and the envelope of HFO signal computed with Hilbert transform. To confirm our hypothesis that HFO is modulated by breathing rhythm we used comodulogram analysis for the two signals. Comodulogram matrix was computed using open-source Python library described in [Dupré la Tour et al., 2017]. We used the “Tort” method from `pactools` Python package, which seems to find a compromise for proper resolution in ‘phase’ and ‘amplitude’ signal and is based on classic phase-amplitude coupling method. We evaluated statistical significance of the coupling using resampling test of MI martices between groups of rats that were under KX and isoflurane anesthesia.

For multielectrode recordings, KX-HFOs significant bursts were detected using 3 standard deviation threshold from top (short silicon probe) and middle (long silicon probe) channels used as a reference. We computed phase shift between channels using maximum correlation score in respect to reference channel and averaged the score across rats. CSDs were reconstructed using kCSD algorithm method from [Potworowski et al., 2012, Chintaluri et al., 2019] and available at https://github.com/Neuroinflab/kCSD-python. We reconstructed CSD first and then filtered spatio-temporal CSD picture in delta 0.3–5 Hz and KX-HFO 80–130 Hz frequency bands. For Multiunit activity analysis we first filtered the signal above 500 Hz for every HFO burst/event. Then we extracted candidates for spikes with 3 standard deviation criterion and represented them as discrete events in time. As a final step we made a histogram from aggregated (across HFO events) spikes and computed Pearson correlation coefficient between spike histogram and average HFO waveform. We repeated this kind of analysis for all channels independently.

Shaded regions in all plots and whiskers of the bar plots represent standard deviation of the mean (s.e.m). For statistical analysis we used one-way ANOVA test for independent experiments or different electrode channels and Student’s paired t-test for different timepoints in a given channel — `f_oneway` and `ttestrel`, respectively, from `SciPy` Python library. All sample groups were tested with Shapiro-Wilk’s test for normality. All data sets passed normality testing and therefore parametric statistics were used. For all figures we used *** for pvalue < 0.001, ** for pvalue < 0.01 and * for pvalue < 0.05 convention. Additionally we used resampling test (100 000 draws with return) to compare modulation index matrices for KX and isoflurane anesthetised rats (n=8).

## 4 Acknowledgments

This work was financed by the National Science Centre (Poland) grant UMO-2016/23/B/NZ/03657 and the Welcome Trust (UK). We would like to acknowledge Prof. Andrzej Wróbel, Piotr Dzwiniel, Anna Posłuszny and Magda Majkowska for assistance with cat experiments.

## 5 Author Contributions

W.S., J.W., E.K. and M.H. conducted the experiments, W.S, D.W., M.W. and M.H. designed the experiments and wrote the paper. W.S and M.H. analyzed the data and designed figures.

## 7 Supplementary figures

**Figure S1.**
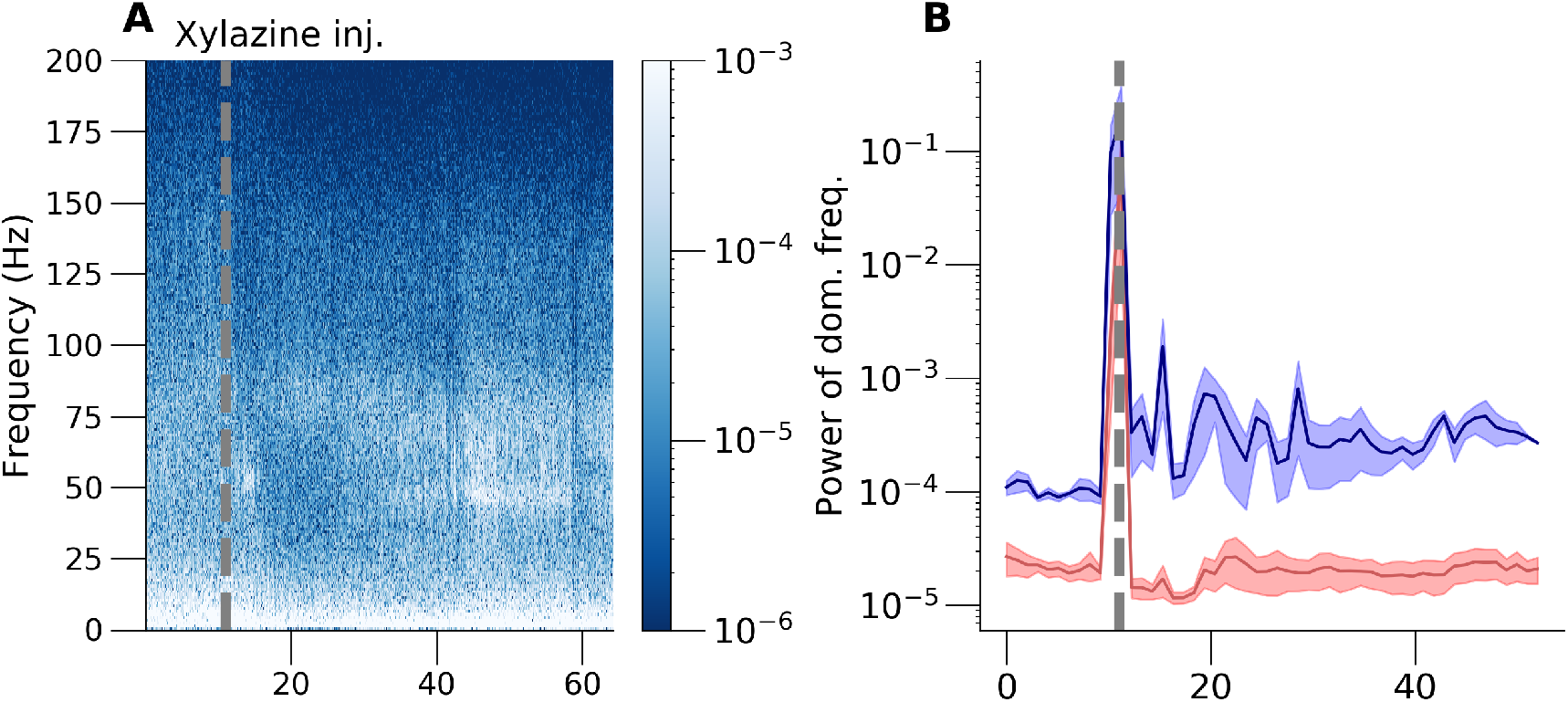
A: Spectrogram of the example rat after xylazine injection. B: Extraction of the power of dominant frequency for Gamma and 80–180 Hz HFO. We did not see any change in HFO after xylazine application alone. C: Spectrogram of the example rat with anesthetic ketamine injection. D: Same type of analysis as B but for ketamine anesthetized rat.

**Figure S2.**
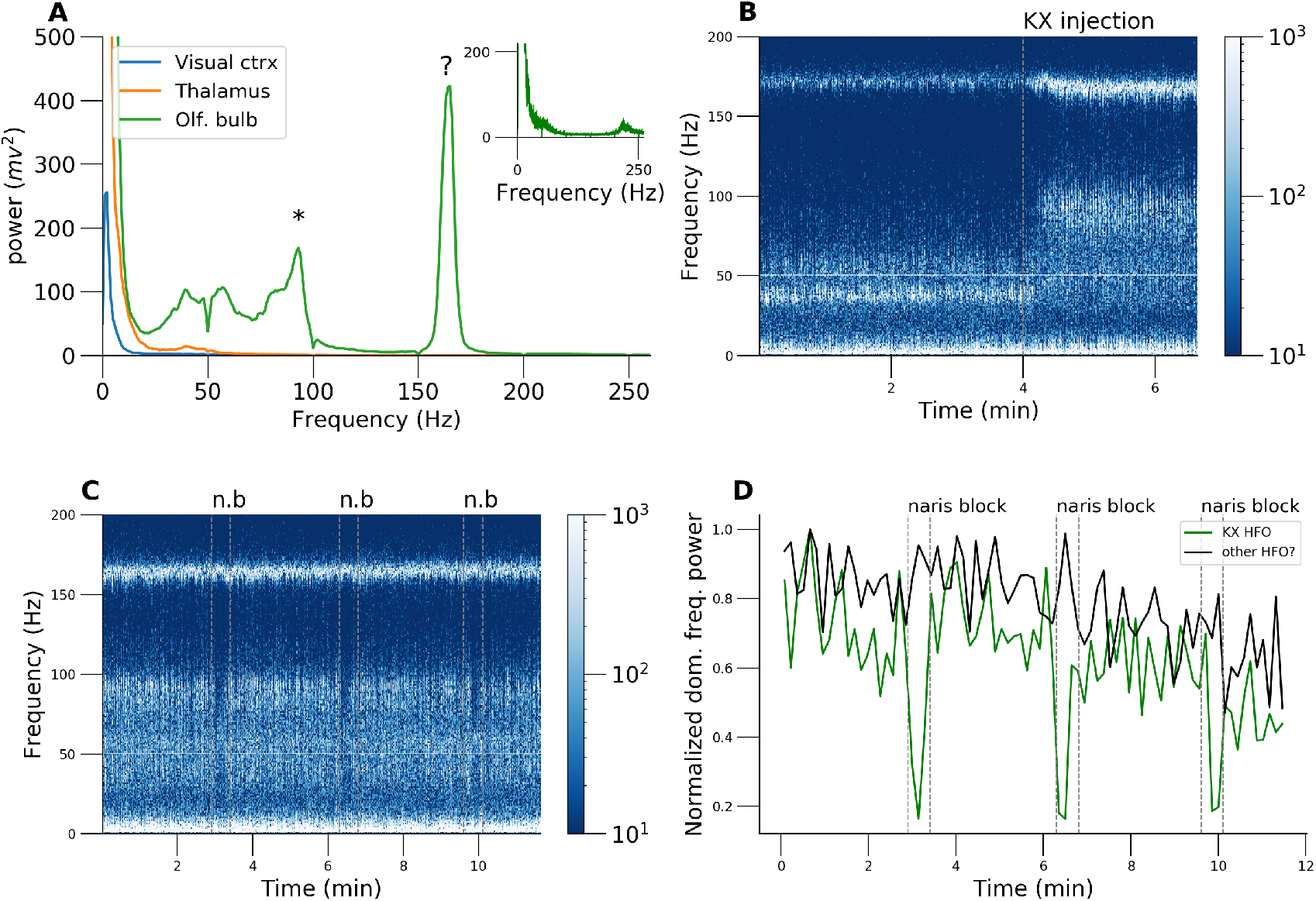
A: Power spectrum of the third cat recorded in OB, LGN and VCx. B: Spectrogram of the response to the renewal of KX anesthesia. C: Spectrogram of the cat under KX anesthesia and naris block experiment. D: Extraction of the power of dominant frequency from 80–130 Hz and 150–170 Hz bands during naris block experiment. Note that 160 Hz oscillation is not reacting to perturbation in respiration.

**Figure S3.**
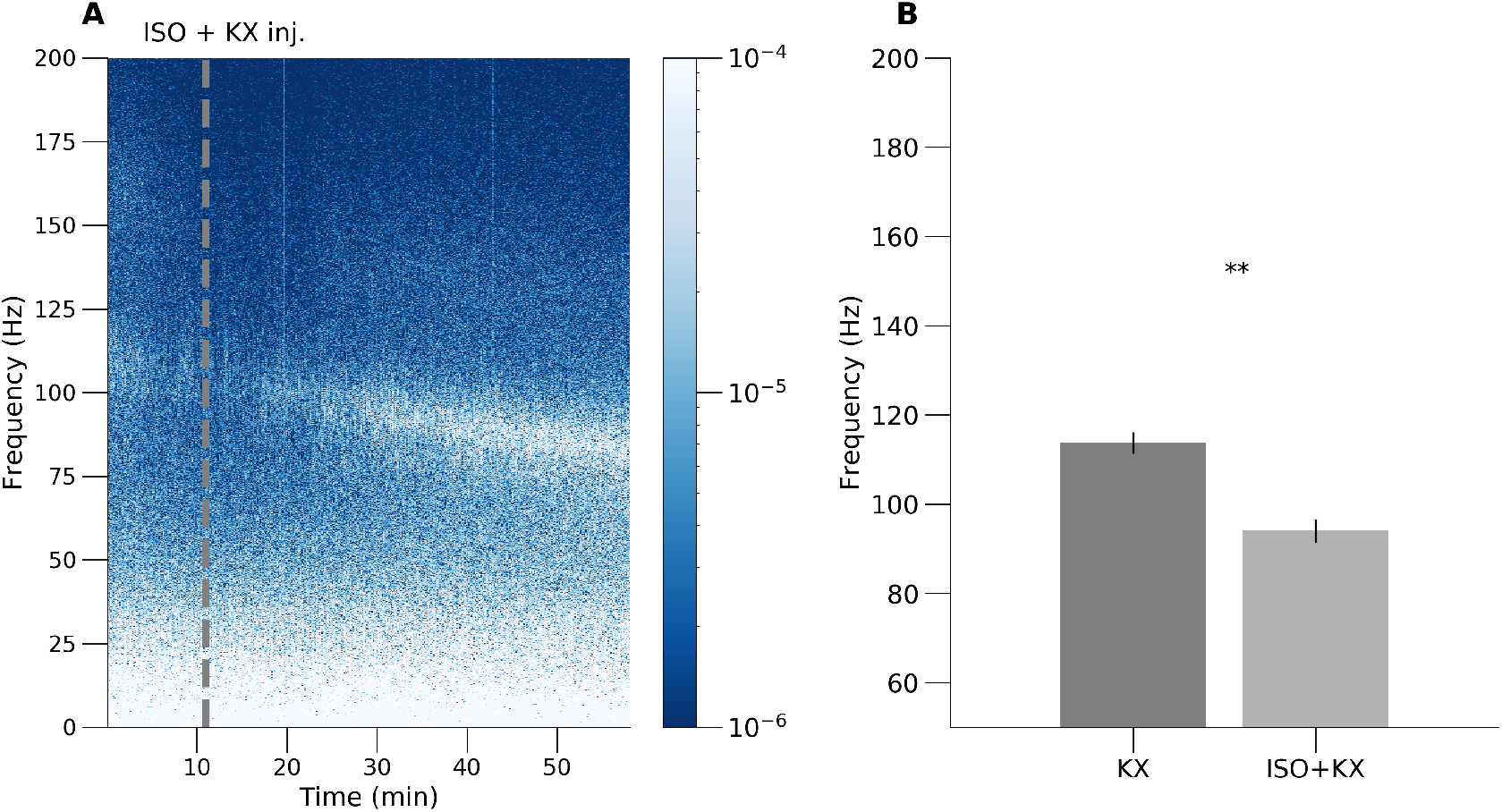
A: Example spectrogram of the chronically implanted rat that was exposed first to isoflurane and then got KX injection. B: Analysis of the frequency reduction in rats that were shortly exposed to isoflurane before KX injection.

**Figure S4.**
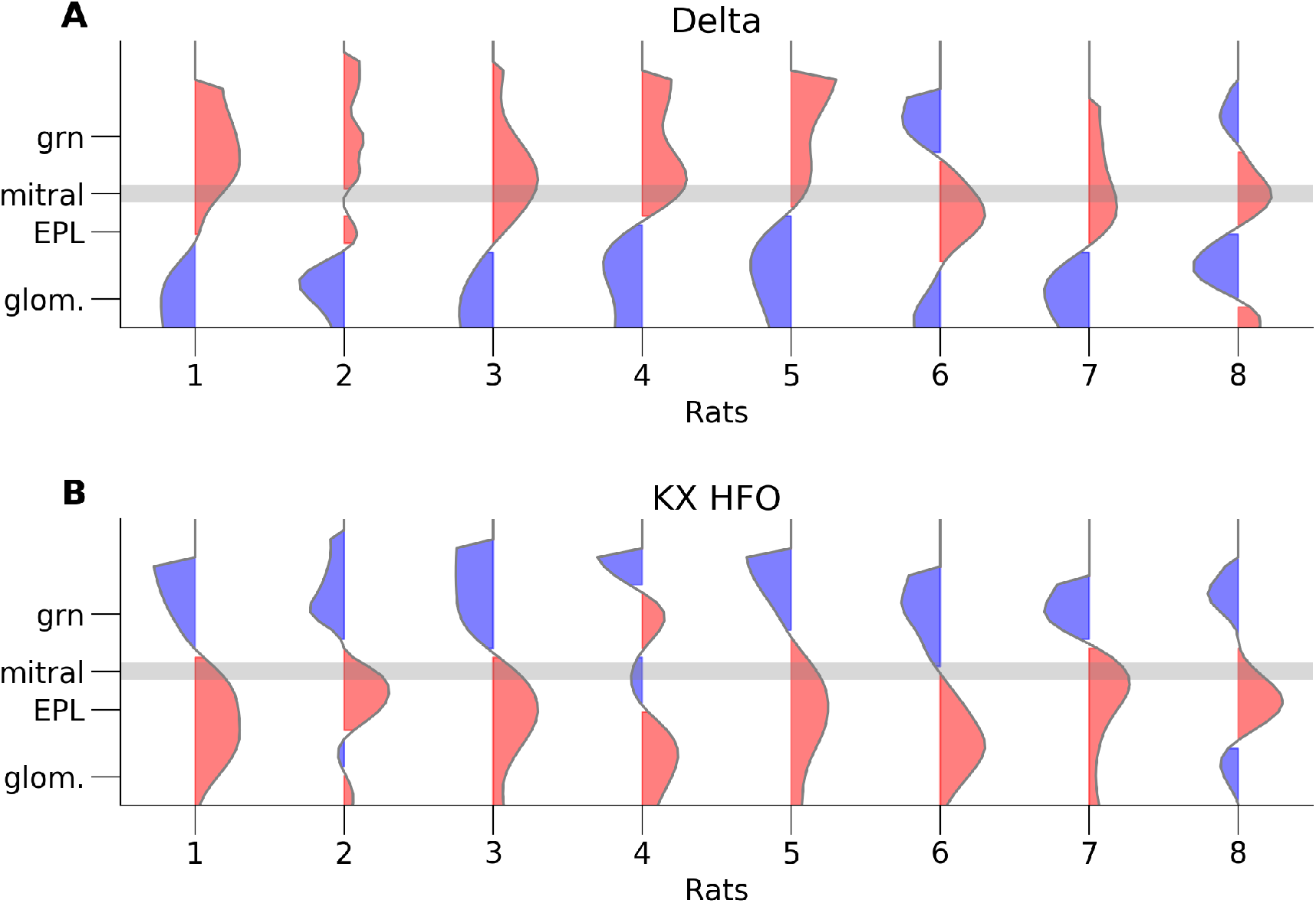
CSD profiles from individual rats 1 to 8 extracted from timepoint zero for delta (A) KX HFO (B) filtered CSDs reconstruction

